# Methylated Cycloalkanes Fuel a Novel Genera in the *Porticoccaceae* Family and Inform Substrate Affinity for a Unique Copper Membrane Monooxygenase

**DOI:** 10.1101/2022.11.07.515388

**Authors:** Eleanor C. Arrington, Jonathan Tarn, Hailie Kittner, Veronika Kivenson, Rachel M. Liu, David L. Valentine

## Abstract

Cycloalkanes are an abundant and toxic class of compounds in subsurface petroleum reservoirs and their fate is quantitatively important to ecosystems impacted by natural oil seeps and spills. In this study, we focus on the microbial metabolism of methylcyclohexane (MCH) and methylcyclopentane (MCP) in the deep Gulf of Mexico. MCH and MCP are often the most abundant cycloalkanes observed in petroleum and a substantial portion of these compounds will dissolve into the water column when introduced at the seafloor via a spill or natural seep. Once dissolved into the water column, the environmental fate of MCH and MCP is presumably controlled by microbial consumption, but little is known about this environmental process. We conducted incubations using fresh Gulf of Mexico (GOM) seawater amended with MCH and MCP at four stations along a transect with a gradient in the influence of natural oil seepage. We observe microbial blooms via optical oxygen sensors that occur at all stations with bloom occurrence among replicate incubations impacted by the proximity of natural seepage. Within all incubations with active respiration of MCH and MCP, we find that *B045*, a novel genus of bacteria belonging to the *Porticoccaceae* family dominates the microbial community. Using seven high-quality metagenome-assembled genomes recovered from microbial blooms on MCH and MCP, we reconstruct the biodegradation pathways and central carbon metabolism of *B045*, identifying a novel clade of the particulate hydrocarbon monooxygenase (*pmo*) that may play a key role in MCH and MCP metabolism. Through comparative analysis of 176 genomes, we parse the taxonomy of the *Porticoccaceae* family and find evidence suggesting the acquisition of *pmo* and other genes related to the degradation of cyclic and branched hydrophobic compounds were likely key events in the ecology and evolution of this group of organisms.

## Introduction

The abundant supply of hydrocarbons from subseafloor reservoirs fuels the global economy and plays a pervasive role in modern society. Marine petroleum inputs, on the order of 1.3 Tg per year, range from catastrophic spills to chronic discharges and impose an array of environmental risks on ecologically sensitive areas [1]. A significant portion of petroleum is a complex mixture of five or six-carbon ring structures with alkyl-substitution, which originate from the degradation of ancient isoprenoid lipids and carotenoid pigments [2]. These compounds have toxicological impacts on aquatic organisms and some pose human health risks via air emissions, a major concern during oil spill response operations and for communities near spill areas [3–5]. Cycloalkanes composed 16% of the total oil and gas released in the Macondo oil from the Deepwater Horizon event, and MCH and MCP represented 2.4% of the total oil released. MCH was the most abundant cycloalkane in Macondo oil [6]. Short-chain cycloalkanes are aqueous soluble, driving their entrapment in the deep ocean when they are introduced at depth via seeps and spills [7]. The long-term fate of petroleum entrapped within the deep ocean is governed by complex hydrocarbon-degrading microbial communities which sequentially consume components of oil and gas substrates [8–13]. To the best of our knowledge, there is no other major source of MCH and MCP in the marine environment, and the microbial communities responsible for their biodegradation are likely specialized to consume petroleum compounds or other similar chemical structures.

Despite their abundance in petroleum, only a few formative studies have focused on cycloalkane metabolism. This work began with Imelik in 1948 with the isolation of the first strain of bacteria (*Pseudomonas aeruginosa*) capable of degrading cyclohexane [14]. Insights from early cultivation studies suggest that in comparison to *n*-alkanes such as pentane, cycloalkanes are more resistant to microbial attack due to additional metabolic steps required for ring cleavage [15, 16]. In cultivated isolates, degradation of substituted cycloalkanes appears to occur more readily than the degradation of unsubstituted forms, particularly if there is an *n*-alkane substitution of adequate chain length for multiple rounds of beta-oxidation prior to ring cleavage [17–19]. MCH and MCP specifically are of interest because their carboxylic acid counterparts (cyclohexane carboxylic acid and cyclopentane carboxylic acid) are often intermediate compounds in beta-oxidized alkylated cycloalkane and alkylated cyclo-carboxylic acid (also called naphthenic acid) degradation [20, 21].

## Materials and Methods

### Incubation Design and Sample Collection

Seawater samples were collected aboard RV Atlantis in June 2015. MCH and MCP incubations were conducted at stations 1 (27° 30.41’ N, 87° 12.41’ W), 2 (27° 15.00’ N, 89° 05.05’ W), 3 (27° 11.60’ N, 90° 41.75’ W) and 4 (27° 38.40’ N, 90° 54.98’ W) with seawater collected from 1,000 m. Seawater collected from the CTD Niskin bottles was transferred to 250 mL glass serum vials using a small length of Tygon tubing. Vials were filled for at least 3 volumes of water to overflow. Care was taken to ensure no bubbles were present before sealing with a polytetrafluoroethylene (PTFE) coated chlorobutyl rubber stopper and crimp cap seal. All bottles, except for unamended blank controls, immediately received 10 μL of MCH or MCP using a gas-tight syringe (Hamilton) and were maintained in the dark at in-situ temperature (4°C). Prior to filling, each serum bottle was fixed with a contactless optical oxygen sensor (Pyroscience, Aachen, Germany) on the inner side with silicone glue, and afterward were cleaned from organic contaminants with rinses of ethanol, 3% hydrogen peroxide, 10% hydrochloric acid, and MilliQ water, and were sterilized via autoclave. Oxygen concentration was monitored approximately every 8 hours with a fiber optic oxygen meter (Pyroscience, Aachen, Germany). Observed changes in oxygen content were normalized to unamended controls to correct for oxygen loss from background respiration processes and variability due to temperature changes. Bloom onset is operationally defined as three consecutive time points with oxygen loss >0.21 μM h^-1^. Importantly, incubations were conducted with no added headspace and were limited to in-situ availability for oxygen, as well as for key nutrients – nitrogen and phosphorous. At the termination of each respiration experiment, samples were harvested and the microbial community was captured on a 0.22-μm polyethersulfone filter.

### DNA extraction, 16S rRNA community analysis

DNA extraction was performed from ¼ of each filter using the PowerSoil DNA extraction kit with the following modifications: 200 μl of bead beating solution was removed at the initial step and replaced with phenol chloroform, the C4 bead binding solution was supplemented with 600 μL of 100% ethanol, and we added an additional column washing step with 650 μL of 100% ethanol. Extracts were purified and concentrated by ethanol precipitation, then stored at −80°C. The V4 region of the 16S rRNA gene was amplified using the method described by [22] with small modifications to the 16Sf and 16Sr primers according to [23, 24]. Amplicon PCR reactions contained 1 μL of template DNA, 2 μL of forward primer, 2 μL of reverse primer, and 17 μL of AccuPrime Pfx SuperMix. Thermocycling conditions consisted of 95° 2 min, 30 cycles of 95°C for 20 secs, 55°C for 15 secs, 72°C for 5 min, and a final elongation at 72°C for 10 min. Sample DNA concentrations were normalized using the SequelPrep Normalization Kit, cleaned using the DNA Clean and Concentrator kit, visualized on an Agilent Tapestation, and quantified using a Qubit Flurometer. Samples were sequenced at the UC Davis Genome Center on the Illumina MiSeq platform with 250nt, paired end reads. A PCR-grade water sample was included in extraction, amplification, and sequencing as negative control to assess for DNA contamination.

Trimmed fastq files were quality filtered using the fastqPairedFilter command within the dada2 R package, version 1.9.3 [25] with following parameters: truncLen=c(190,190), maxN=0, maxEE=c(2,2), truncQ=2, rm.phix=TRUE, compress=TRUE, multithread=TRUE. Quality filtered reads were dereplicated using derepFastq command. Paired dereplicated fastq files were joined using the mergePairs function with the default parameters. A Single Nucleotide Variant (SNV) table was constructed with the makeSequenceTable command and potential chimeras were removed denovo using removeBimeraDenovo. Taxonomic assignment of the sequences was done with the assignTaxonomy command using the Silva taxonomic training dataset formatted for DADA2 v132 [26, 27]. If sequences were not assigned, they were left as NA.

### Metagenomic Reconstruction

Metagenomic library preparation and shotgun sequencing were conducted at the University of California Davis DNA Technologies Core. DNA was sequenced on the Illumina HiSeq4000 platform, producing 150-base pair (bp) paired-end reads with a targeted insert size of 400 bp. Quality control and adaptor removal were performed with Trimmomatic [28] (v.0.36; parameters: leading 10, trailing 10, sliding window of 4, quality score of 25, minimum length 151 bp) and Sickle [29] (v.1.33 with paired-end and Sanger parameters). The trimmed high-quality reads were assembled using metaSPAdes [30] (v.3.8.1; parameters *k* = 21, 33, 55, 77, 88, 127). The quality of assemblies was determined using QUAST [31] (v.5.0.2 with default parameters). Sequencing coverage was determined for each assembled scaffold by mapping high-quality reads to the assembly using Bowtie2 [32](v.2.3.4.1; default parameters) with Samtools [33] (v.1.7). Contigs greater than 2,500 bp were manually binned using Anvi’o with Centrifuge [34, 35] (v.1.0.1) based on coverage uniformity (v.5). Quality metrics for metagenome-assembled genomes (MAGs) were determined using CheckM [36] (v.1.0.7; default parameters). The taxonomy of each MAG was classified using GTDB-Tk (v.1.0.2) against The Genome Taxonomy Database [37] (https://data.ace.uq.edu.au/public/gtdb/data/releases/release89/89.0/, v.r89).

### Metagenome Annotation

Open reading frames were predicted for MAGs using Prodigal [38] (v.2.6.3; default parameters). Functional annotation was determined using HMMER3 [39](v.3.1b2) against the Pfam database (v.31.0) with an expected value (e-value) cutoff of 1 × 10^-7^ and KofamScan (v.1.1.0) [40] against the Hidden Markov model (HMM) profiles for Kyoto Encyclopedia of Genes and Genomes and Kegg Orthology (KEGG/KO) with a score cutoff of 1 × 10^-7^. To find hits for *almA* we used Pfam (<u>PF00743</u>), for *rhdA* we used Pfam (<u>PF00848</u>) and for *pHMO* we summed KO hits (subunit a: K10944; subunit b: K10945; subunit c: K10946). For alkane-1-monooxygenase (*alkB*) detection we used KofamScan with <u>K00496</u> to search for *alkB*.

### Phylogenetics

To define genome phylogenomic relationships of MAGs, 16 universal ribosomal proteins (RPs) were used (L2-L6, L14-L16), L18, L22, L24, S3, S8, S10, S17, and S19. For phylogenies of metabolic genes as well ribosomal proteins, all representative sequences and concatenated alignments that contained <25% informative sites were excluded in tree construction. Each protein was aligned using MUSCLE (v.3.8.425) [41]. All columns with >95% gaps were removed using TrimAL [42]. Maximum-likelihood phylogenetic analysis of concatenated alignment was inferred by RAxML [43] (v.8..9; parameters: raxmlHPC -T 4 -s input -N autoMRE -n result -f a -p 12345 -x 12345 -m PROTCATLG). The resulting trees were visualized using FigTree [44] (v.1.4.3).

## Results

### Methylated Cycloalkane Bloom Occurrence

In this study, we conducted incubations at-sea with freshly-collected water from the deep ocean (1,000 m) along a transect within the Gulf of Mexico (GOM) (Fig. 1). MCH and MCP metabolism was observed through a closed-system optical oxygen technique in which population blooms were defined by the exponential rise in respiration at > 0.21 μM per hour. At all stations within the Gulf of Mexico blooms occurred at 18-21 days. However, bloom frequency among incubations varied in the seep-replete [45] Northwest Gulf of Mexico compared to the seep-depleted region located to the Northeast. At stations, #2, #3, and #4 over 50% of the incubations bloomed within 21 days for both MCH and MCP. More specifically, for MCH: station #1 located far from seepage saw 3 of 11 incubations bloom within 30 days; at station #2, 4 of 6 bloomed, at station #3, 4 of 6 bloomed; and at station #4 located near to seepage, 9 of 11 bloomed. For MCP a similar pattern is observed: station #1 (0 of 11 bloomed), station #2 (4 of 6 bloomed), station #3 (4 of 6 bloomed), and station #4 (10 of 11 bloomed). The average respiration profile for blooms on MCH and MCP also showed similar patterns in oxygen loss with time across each station (Fig. 2).

**Figure 1.**
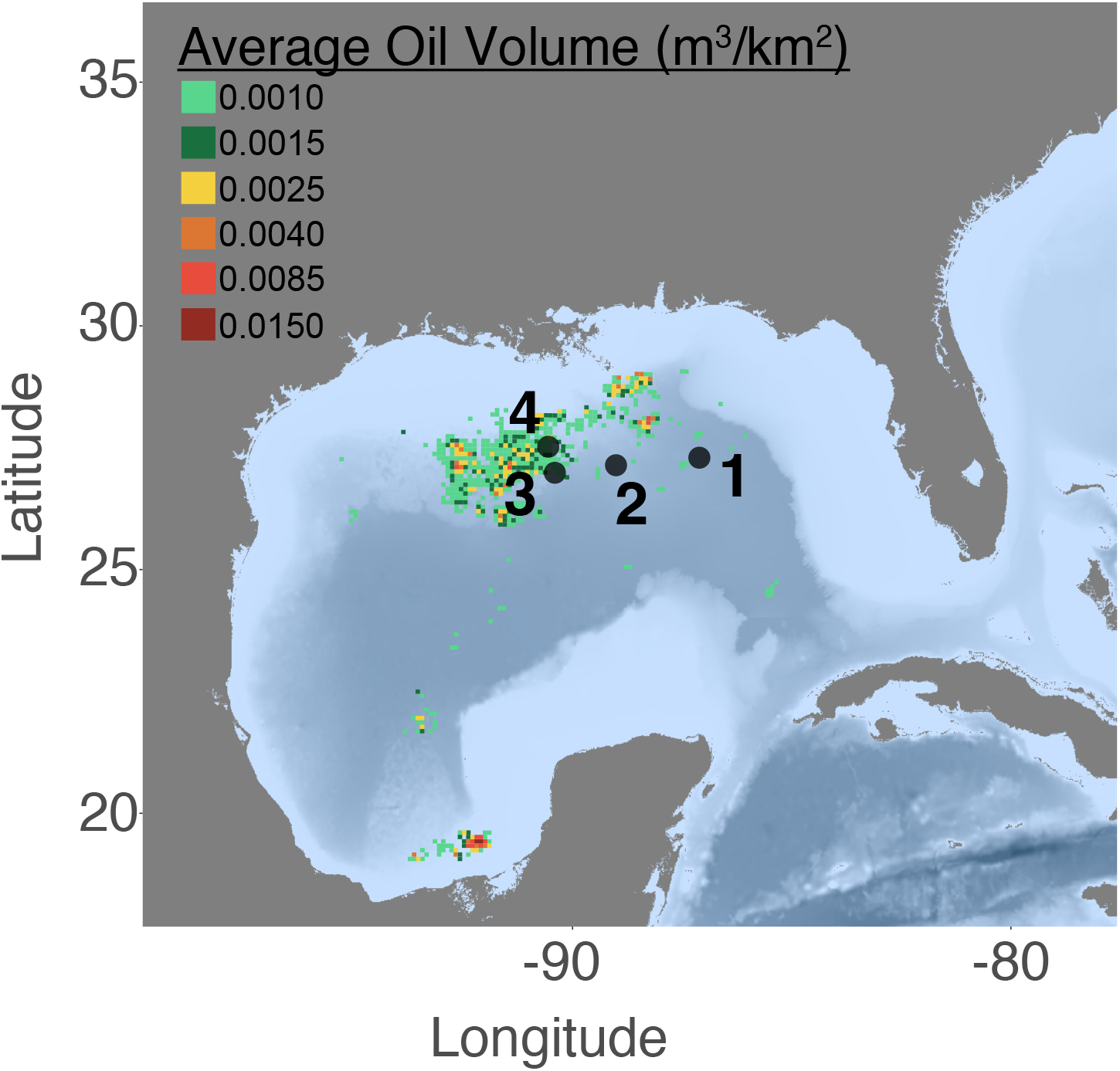
Sampling stations relative to natural oil seepage in the Gulf of Mexico.

**Figure 2.**
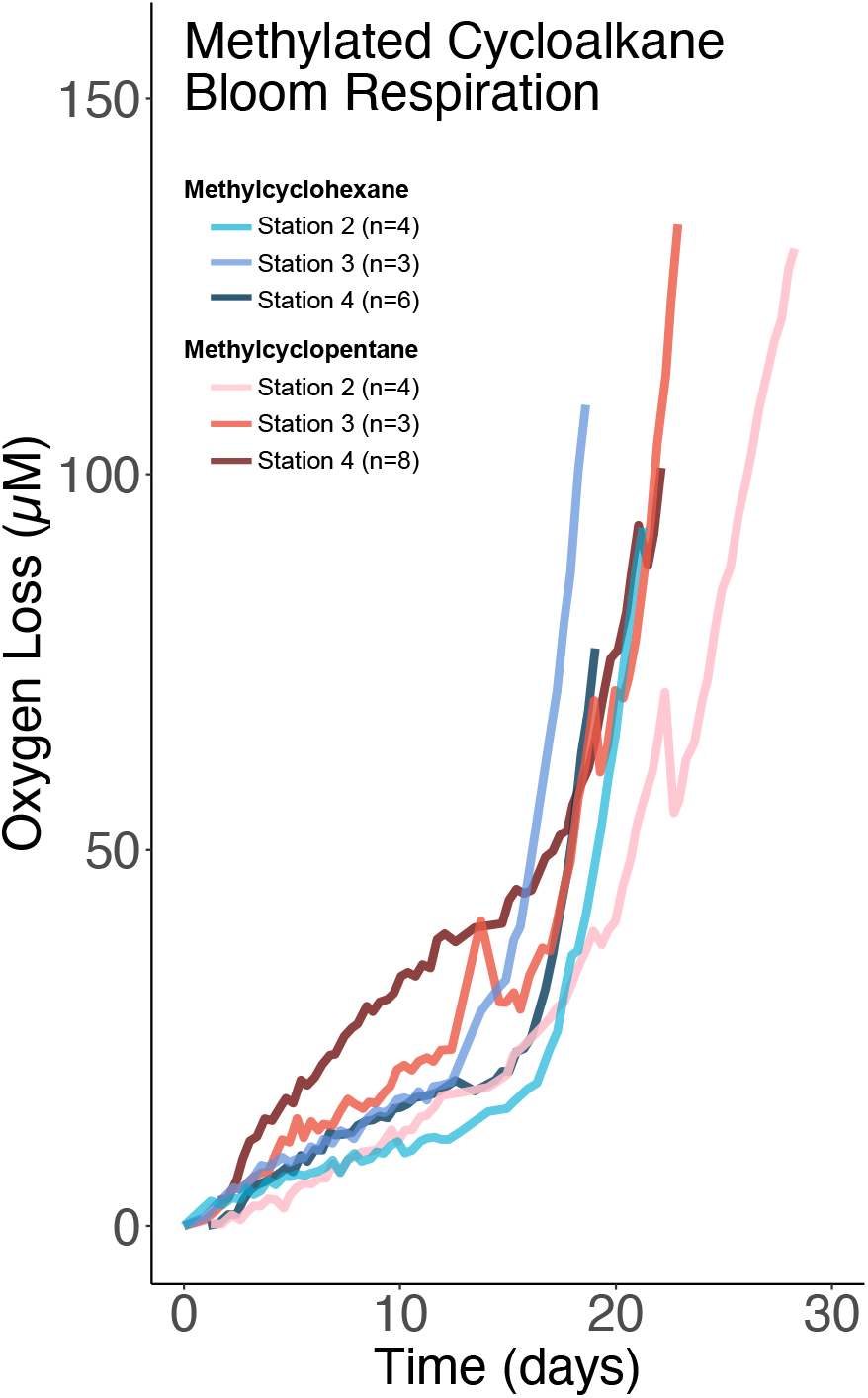
Average oxygen loss with time for blooms (defined by oxygen loss >0.21 μM h^-1^) on methylcyclopentane and methylcyclohexane at station 2, 3, and 4.

### Microbial Community Response to Cycloalkanes

MCH and MCP enrichments typically elicited the emergence of a single dominant taxa comprising >50% of the 16S rRNA amplicon sequences across all stations (Fig. 3). The full taxonomy of the closest match to the dominant SNV (observed in all but two samples regardless of bloom status) is as follows in the Silva and NCBI database: Bacteria; Proteobacteria; Gammaproteobacteria; *Cellvibrionales; Porticoccaceae; C1-B045* [27]. Blooms containing abundant taxa other than *C1-B045* occurred four times out of the twenty-two (MCH- and MCP-derived) blooms investigated (Fig 3). These other blooming taxa belong to the genera *Cycloclasticus, Colwellia*, and *Moritellaceae* (each >30% abundant in a single bloom). Only one bloom at station 1 with *Cycloclasticus* occurred without the co-occurrence of *C1-B045*. The *C1-B045* SNV is a novel (uncultivated) bacteria at the genera level that was first detected in deepsea hydrothermal vents [46], has been detected in other oil-contaminated regions [47], and is distantly related to SAR92 [48]. To the best of our knowledge, a complete phylogenomic analysis of the *Porticoccaceae* family is not presently available.

**Figure 3.**
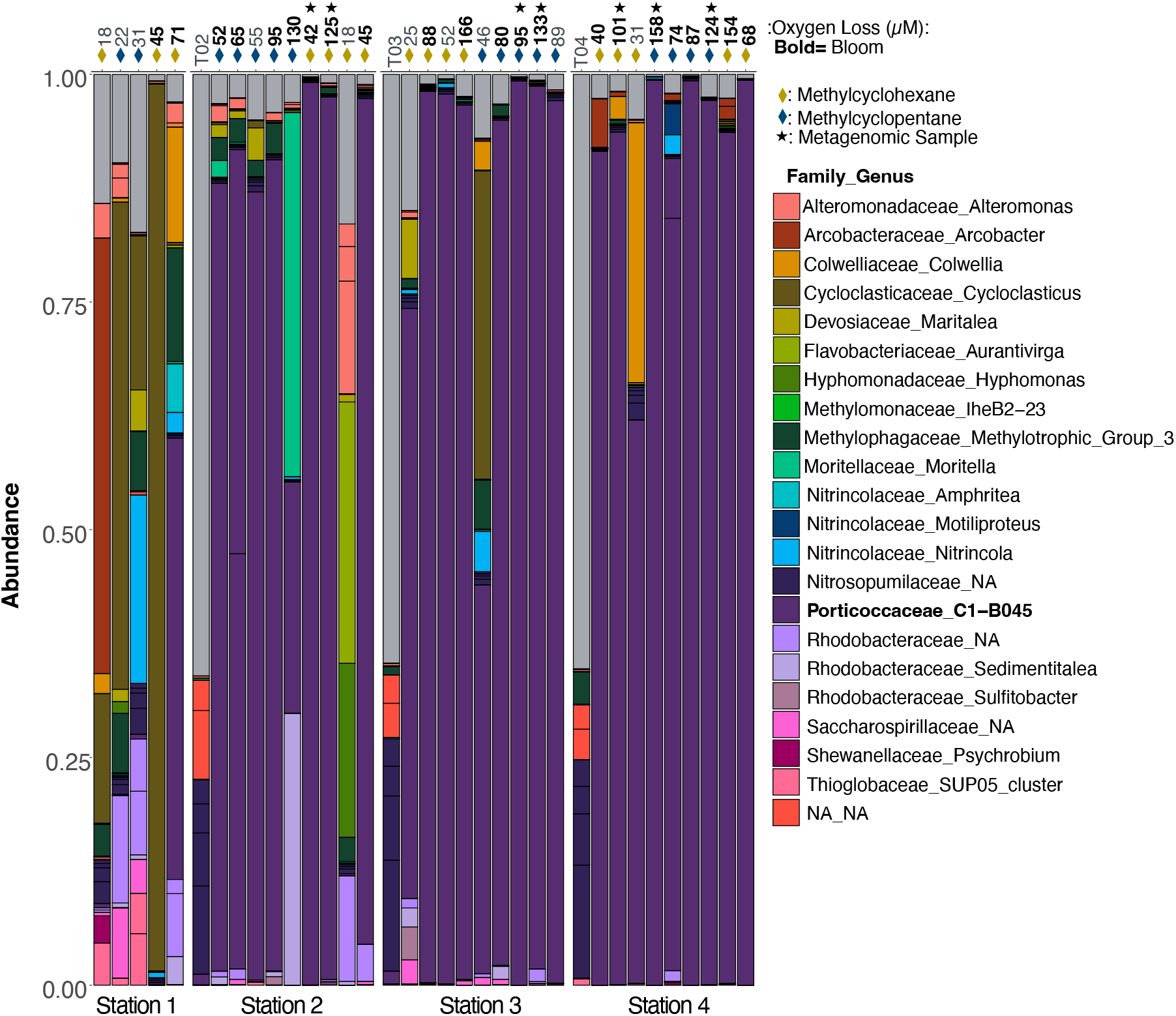
Microbial community composition of methylcyclohexane (MCH) and methylcyclopentane (MCP) blooms informed via the V4 region of the 16S rRNA gene. Initial environmental samples (denoted “T0”). The amount of oxygen loss in the incubation when DNA collection occurred is denoted by the values at the top of the figure. Bold oxygen loss values indicate incubation met the oxygen loss criteria for a bloom. “NA_NA” indicates no taxonomic representative at the family or genera level. Diamond color indicates which compound was added (MCP or MCH). Black stars note which samples were sequenced further for metagenomic analyses.

### Genome Reconstruction and Taxonomic Inferences

We reconstructed high-quality metagenomes from seven MCH and MCP treatments in a state of active respiratory bloom, with completeness >97% and redundancy <2% (black stars in Fig. 3, Table 1). Two MAGs, named “MCH_2_108” and “MCH_2_109”, originated from station #2 (further from natural seepage) with MCH enrichment; two MAGs, named “MCP_3_146” and “MCP_3_148”, originated from MCP enrichments at station #3; and three MAGs, named “MCP_4_184”, “MCP_4_160”, and “MCH_4_158” originated from MCH and MCP enrichment in the natural seep region at station #4. Each MAG was recovered from biologically independent incubations, yet every major component of metabolism and every taxonomic marker was nearly identical across each; therefore, we will refer to them as “B045-MAG”. Using our high-quality genomes as queries to the Genome Taxonomy Database we found the family *Porticoccaceae* has recently been reclassified to resolve polyphyletic phylogenetic relationships among the *Cellvibrionales* family [49]. The taxonomy of the B045-MAG according to GTDB-Tk is Bacteria; Proteobacteria; Gammaproteobacteria; Pseudomonadales; *Porticoccaceae; 500-400-T64*. The genera designation “500-400-T64” refers to a non-standard placeholder name assigned by GTDB-Tk. There have been no genus names proposed formally for this group of genomes, so for the purpose of this study, we refer to the genus as *B045*.

**Table 1.** Bin statistics for metagenomic constructions from this study. Completion and redundancy were calculated with CheckM.

| MAG Name | Number of Contigs | Genome Size | Main genome contig N/L50: | Completion | Redundancy |
| --- | --- | --- | --- | --- | --- |
| MCH_2_108 | 4 | 3.156 MB | 2/832.349 KB | 100.0% | 0.5% |
| MCH_2_109 | 3 | 3.118 MB | 2/1.058 MB | 99.8% | 0.5% |
| MCP_3_148 | 4 | 3.141 MB | 2/1.167 MB | 99.8% | 0.5% |
| MCP_3_146 | 4 | 3.159 MB | 1/1.823 MB | 99.8% | 0.6% |
| MCP_4_184 | 5 | 3.113 MB | 2/732.488 KB | 99.8% | 0.5% |
| MCP_4_160 | 11 | 3.115 MB | 2/579.437 KB | 99.8% | 0.5% |
| MCH_4_158 | 156 | 2.762 MB | 24/36.15 KB | 96.0% | 0.5% |

## Discussion

### Methylcyclohexane Metabolism

Within the B045-MAG we found the potential for cycloalkane utilization in both catabolism and anabolism. B045 also appears to metabolize both MCH and MCP, however, given there is a large amount of redundancy between the two pathways and the larger number of studies conducted on cyclohexane metabolism in cultured isolates, we mainly focus here on MCH metabolism. Two different biodegradation strategies for the compound cyclohexane have been identified in bacteria; a lactone formation pathway and an aromatization pathway. The intermediate compounds for the aromatization pathway were observed by [50] in *Rodococcus* sp. EC1 whereby cyclohexane was converted to phenol. The lactone formation pathway was characterized by Stirling et al. 1977 in *Nocardia* finding that growth, respiration, and enzymes from culture extracts are consistent with cyclohexane degradation via cyclohexanol, cyclohexanone, caprolactone, and *ε*-hydroxycaproate [17]. This metabolic route for cyclohexane consumption was further corroborated in *Pseudomonas* and *Xanthobacter* cultures [51, 52]. Enzymes used in this pathway include cyclohexanol dehydrogenase, cyclohexanone monooxygenase, and *ε* caprolactone hydrolase. The cyclopentane degradation pathway, first elucidated by [53], occurs via cyclopentanol, cyclopentanone, 5-valerolactone, 5-hydroxyvalerate, glutarate, and acetyl-CoA [53]. This pathway has been observed in *Comamonas* sp. and *Acinetobacter* with significant efforts to define key enzymes including cyclopentane monooxygenase, cyclopentanol dehydrogenase, cyclopentanone 1,2-monooxygenase, a ring-opening 5-valerolactone hydrolase, 5-hydroxyvalerate dehydrogenase, and 5-oxovalerate dehydrogenase [54]. To the best of our knowledge, no studies have found aromatic intermediates during MCH degradation and furthermore, the phenol biodegradation pathway is absent from B045-MAG, therefore we eliminated aromatization from our hypothesized MCH pathway.

MCH consumption with a lactone intermediate involves the oxidative attack from within the cyclic ring, causing the formation of the following structures: a cyclic alcohol, a ketone, then a lactone, followed by conversion to a carboxylic acid that is beta-oxidized and shunted into central carbon metabolism. A number of studies have found evidence for this general pathway in MCH consumption, however, two studies found differences as to which carbon relative to the methyl group is initially oxidized. Tonge and Higgins found that the isolate *Norcardia petroleophilia* grows on methylcyclohexane, and found 3-methylcyclohexanol and 3-methylhexanone in the growth medium during exponential growth [55]. Koma et al., 2004 found 4-methylcyclohexanol, 4-methylcyclohexanone and methyl-ε-caprolactone as intermediates of MCH degradation by *Rhodococcus* [56].

MCH differs from cyclohexane metabolism in that there could be a third form of metabolism, whereby the methyl group is first oxidized, forming cyclohexymethanol, then cyclohexanecarboxylic acid, which is converted over multiple metabolic steps to pimeloyl-CoA and downstream is beta-oxidized and shunted into the tricarboxylic acid cycle. This pathway for methyl group oxidation shares a number of metabolic steps with benzoate metabolism. Lloyd-Jones and Trudgill studied a bacterial consortium of *Rhodococcus, Flavobacterium*, and *Pseudomonas spp*. isolated with methylcyclohexane as the sole carbon source from contaminated soil and water samples from an oil refinery. This bacterial consortium showed catabolic flexibility (within ring and methyl group oxidation) based on the substrates it could grow on, which included cyclohexanecarboxylic acid, as well as 2-methylcyclohexanol, 3-methylcyclohexanol, and 4-methylcyclohexanol indicating some organisms may be able to oxidize MCH multiple ways [57]. These seminal works on cycloalkane enzymology are vital points of comparison for our study. We have outlined two metabolic pathways and the associated genes for initial oxidation within the cyclic ring to 3-methylcyclohexanol and as well as initial oxidation of the methyl group (Fig. 4, Fig S1, Table 2, Table S1). B045-MAG is missing 3 genes out of 15 for the pathway involving methyl-group oxidation whereas it contains the complete set of genes for the initial oxidative attack to 3-methylcyclohexanol. We hypothesize that MCH is oxidized via 3-methylcyclohexanol but recognize that it is possible the missing steps for oxidation of the methyl group could be too divergent to be recognized by our annotation approach or B045-MAG could encode novel enzymes to perform these steps. It is also possible that B045-MAG could utilize multiple metabolic strategies contemporaneously.

**Figure 4.**
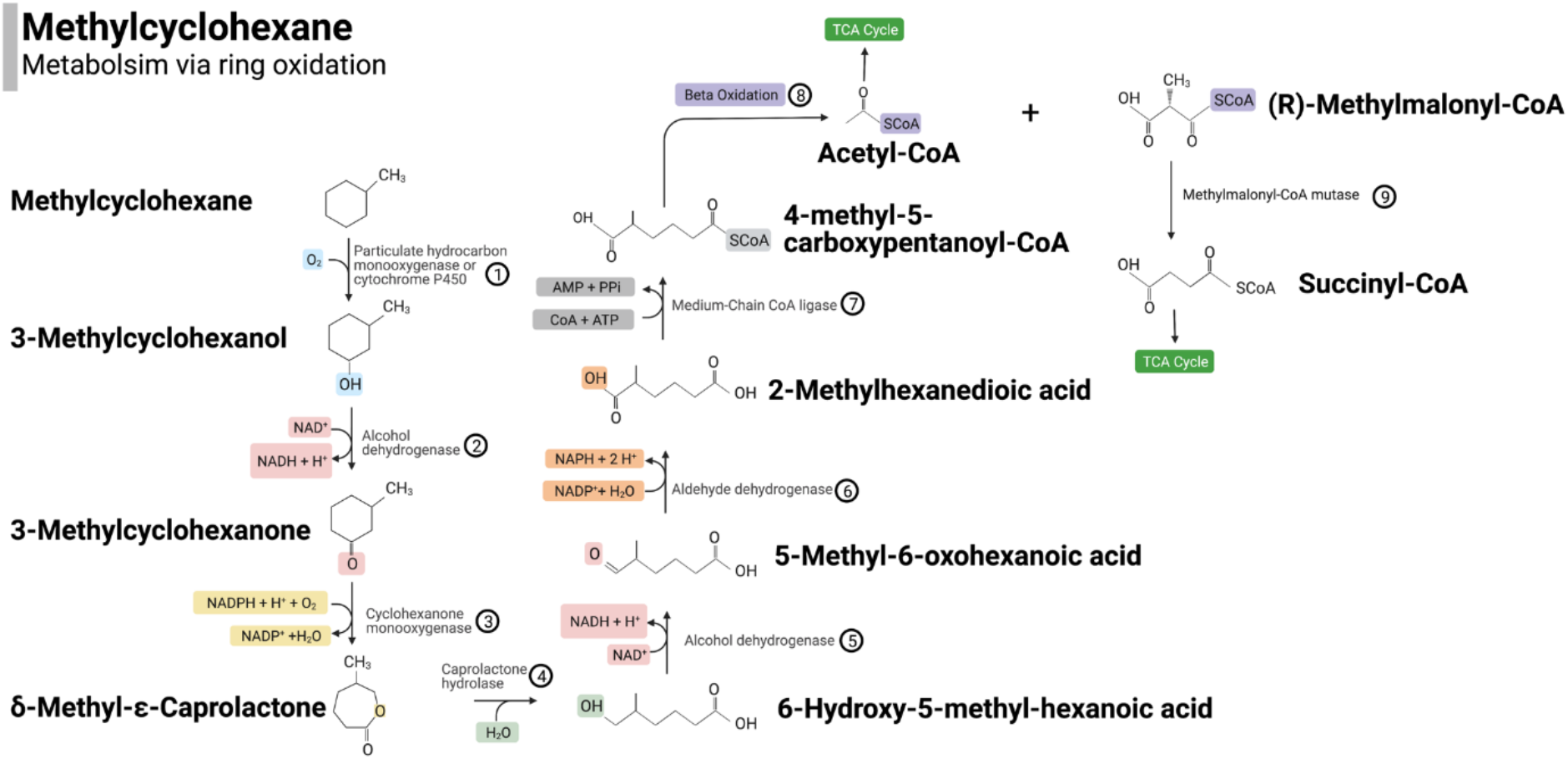
Hypothetical pathway for methylcyclohexane consumption via ring oxidation. Enzymes are denoted by each number with a circle around it. B045-MAG encodes all genes in step 1-9 as well as the tricarboxylic acid cycle. Numbers correspond to enzymes and are elaborated on in Table 2.

**Table 2.** KEGG Gene Identifiers used for Fig. 4.

| Metabolic Step | Enzyme Name | KO/KEGG ID |
| --- | --- | --- |
| ① | Particulate Hydrocarbon Monooxygenase or Cytochrome P450 | K10944 or K15486 |
| ② | Cyclohexanol Dehydrogenase | K19960 |
| ③ | Cyclohexanone Monooxygenase | K00499 |
| ④ | Gluconolactonase | K03379 |
|  | Epsilon-Lactone Hydrolase | K14731 |
| ⑤ | Alcohol Dehydrogenase | K13954 |
| ⑥ | Aldehyde Dehydrogenase | K00138 |
| ⑦ | Medium-Chain CoA Ligase | K23617 |
| ⑧ | Acyl-CoA Dehydrogenase | K00249 |
|  | Enoyl-CoA hydratase | K01692 |
|  | 3-Hydroxyacyl-CoA Dehydrogenase | K00022 |
| ⑨ | Methylmalonyl-CoA mutase | K01848, K01849 |

### Initial Oxidation of MCH and MCP

The first step in hydrocarbon consumption is oxidation. For methylated cycloalkanes initial oxidative attack can be performed by a number of enzymes including the soluble non-heme iron monooxygenase (sMMO) encoded by *mmoABC*, membrane-bound heme-containing cytochrome P450 enzymes of the CYP153 family (CYP), non-heme iron monooxygenase (ALK) encoded by *alkB*, flavin-binding monooxygenases (ALM) encoded by *almA*, or the particulate copper-containing monooxygenases pHMO encoded by *pmoCAB* [58]. The pHMO enzyme belongs to a larger family of enzymes known as the copper membrane monooxygenases (CuMMOs) which contain three different primary substrates: methane for particulate methane monooxygenase (pMMO), short chain alkanes and alkenes for the particulate hydrocarbon monooxygenase (pHMO), and ammonia for ammonia monooxygenase (AMO). CuMMOs have wide substrate affinity including ammonia, methane, ethane, ethylene, propane, butane, and acetone [59–63].

We searched for enzymes to catalyze the initial oxidation of methylcyclohexane and found potential matches for the enzymes ALK, pHMO, CYP, and ALM. ALM enzymes encoded by *almA* has been experimentally validated to function on long-chain alkanes (*n*C_18_-*n*C_36_) and branched alkanes (pristane), yet to our knowledge has never been implicated in alkane consumption <18 carbons in length; therefore it is unlikely to initially oxidize MCH and MCP [64–66]. The *alkB-like* homolog contained only one of eight essential histidine residues and formed a monophyletic clade with the ancestrally related enzyme for bacterial and eukaryotic fatty acid desaturase, therefore, we find it unlikely to have activity on MCH (data not shown) [67]. We found a single copy of a gene belonging to the cytochrome P450 enzyme family within B045-MAG, and we do not exclude the possibility that this gene could oxidize cycloalkanes, although we note the phylogenetic relationships among CYP enzymes are often difficult to resolve and non-functional forms of this enzyme have been identified in alkane degrading bacteria [68]. All three subunits (*pmoCAB*) for the pHMO enzyme are encoded within the B045-MAG. This enzyme likely has a role in hydrocarbon oxidation in the B045, possibly on MCH and MCP, given the number of genes B045-MAG encodes for catabolism and anabolism of hydrocarbon substrates (Fig.4, Fig. S1, Table 2, Table S2). For the purposes of our metabolic reconstruction, we hypothesize that the initial oxidation of MCH and MCP is performed by either CYP or pHMO (Fig. 4 and Fig S1).

### Alcohol Conversion to TCA Cycle

For the second step of MCH consumption, the B045-MAG utilizes an alcohol dehydrogenase to convert 3-methylcyclohexanol to 3-methylcyclohexanone. For the third step in MCH consumption, 3-methylcyclohexanone is transformed to delta-methyl-epsilon caprolactone using a cyclohexanone monooxygenase that is evolutionarily related to *almA* and catalyzes Baeyer-Villiger oxidations on a number of ketones including cyclopentanones and cyclobutanones [69]. We found 7 copies of cyclohexanone monooxygenase genes present in B045-MAG, possibly illustrating the diversity of cyclic compounds targeted by the organism. In the fourth step of methylcyclohexane consumption, the delta-methyl-epsilon-caprolactone is hydrolyzed to form 6-hydroxy-5-methyl-hexanoic acid, which is a function likely performed by the epsilon-lactone hydrolase in the B045-MAG. Another alcohol dehydrogenase is used in the fifth step of consumption to convert the 6-hydroxy-5-methyl-hexanoic acid to 5-methy adipate semialdehyde. The sixth step of consumption requires an aldehyde dehydrogenase to convert 5-methyl adipate semialdehyde to 5-methyl adipate. In the seventh step, medium-chain CoA ligase prepares the compound for beta-oxidation by adding CoA resulting in 5-methyl adipyl CoA. Beta-oxidation subsequently occurs with acetyl-CoA and methyl malonyl CoA as the products. Interestingly, the B045-MAG contains many copies of the genes required for beta-oxidation, including 17 individual copies of acyl-CoA dehydrogenase per genome, leading us to suggest that B045-MAG may specialize in beta-oxidation of a variety of substrates. The remaining methyl malonyl CoA contains a methyl group in the beta-carbon position preventing traditional beta-oxidation. The methyl malonyl CoA mutase is used in the final step to overcome this obstacle and convert methyl malonyl CoA to succinyl-CoA. Both the acetyl-CoA and succinyl-CoA resulting from the metabolic pathway are shunted into central carbon metabolism via the tricarboxylic acid cycle (TCA).

### Phylogenomic Analysis of B045-MAG

To gain an understanding of how hydrocarbon metabolic capability within the *Porticoccaceae* family relates to ecological and evolutionary patterns, we compiled a total of 176 high-quality (>70% complete, <5% redundant) *Porticoccaceae* genomes from a variety of environments using publicly available data. These environments included marine oligotrophic surface waters, marine mesopelagic waters, coastal sand, hydrothermal vents, marine phytoplankton blooms at a variety of latitudes, contaminated groundwater, and oil mesocosms from a variety of sources including Labrador Sediment, Gulf of Mexico, and the Douglas Channel. Each genome was analyzed using the Genome Taxonomy Database (GTDB-Tk) [37] (v.1.0.2) which uses 120 bacterial marker genes to classify the taxonomic identity of each genome and then uses a threshold of average nucleotide identity >95% to designate microbial species. The 176 genomes were used to construct a phylogenomic tree of the *Porticoccaceae* family and each genus was labeled according to designations provided by GTDB-Tk (Fig. 5). For all genera without a cultivated isolate, GTDB-Tk provides a “non-standard placeholder name” which is typically a MAG “isolate name” from the National Center Biotechnology Information (NCBI) database of a genome within that clade (Fig. 5).

**Figure 5.**
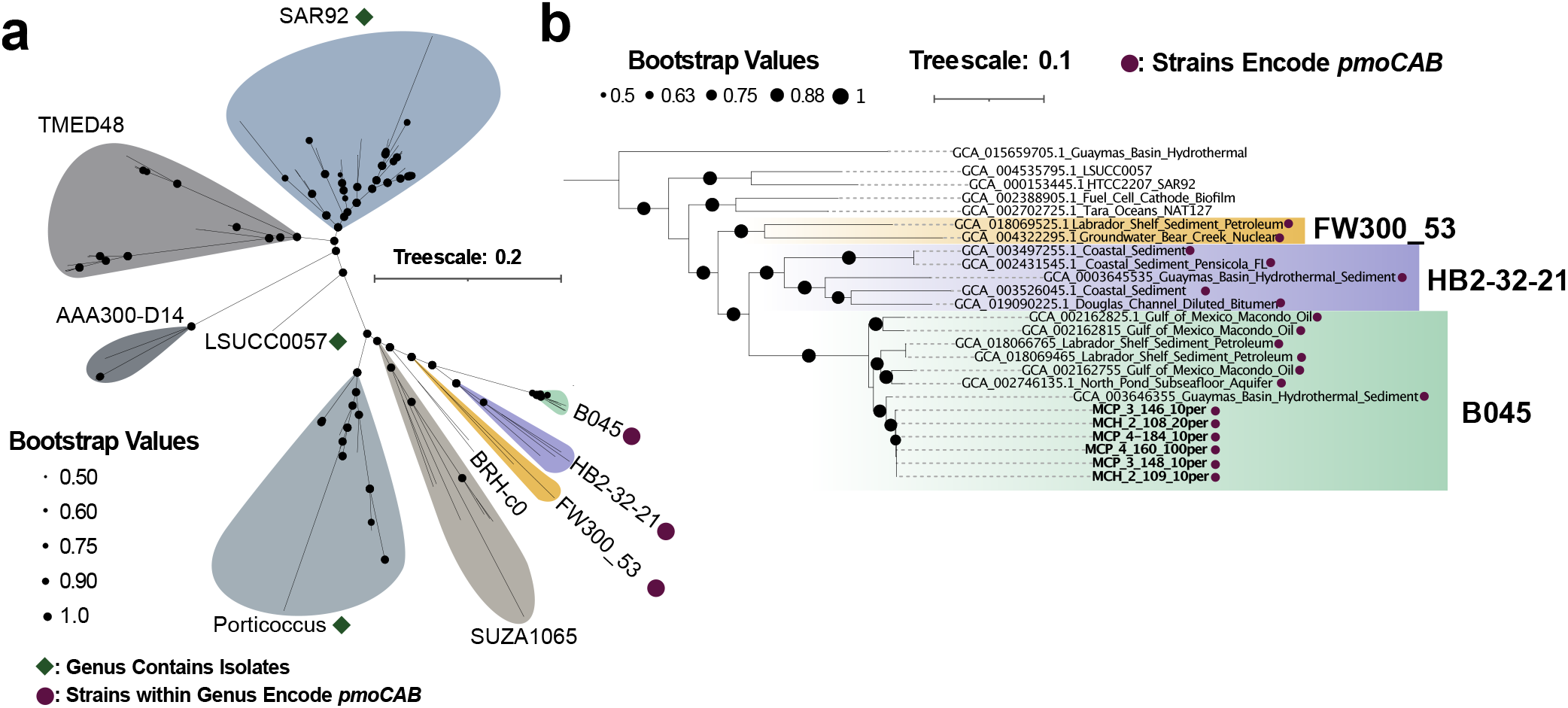
Phylogenomic analysis of *Porticoccaceae* ribosomal proteins. **a** Phylogeny constructed using 16 ribosomal proteins, colored subclades are designated and labeled based on GTDB-Tk classification of genera. Cultivated isolates are noted with green diamond. Genera that encode *pmo*CAB are highlighted with maroon circle. **b** Phylogenomic tree of B045-MAG and close relative ribosomal proteins rooted on SAR92. Genomes that encode *pmo*CAB are noted with maroon circle. The tree scale bar represents substitutions per site.

The B045-MAG forms a well-supported monophyletic clade with other genomes classified by GTDB-Tk as the same genus. Genomes within the *B045* genus all appear to exist in environments where petroleum compounds are present including two genomes from Gulf of Mexico enrichments with Macondo oil, two genomes from Labrador Shelf sediment inoculated with diesel, a single genome from the Guaymas Basin, and a single genome from the North Pond subseafloor aquifer. Both Guaymas Basin and North Pond sites are locations where petroleum compounds have been noted in previous studies [70, 71]. The most closely related genera to *B045* are the *HB2-32-21* and *FW300_53*. The *HB2-32-21* include genomes from the Douglas Channel incubated with diluted bitumen, coastal sediment from Pensacola, Florida, and another genome from a Guaymas Basin hydrothermal vent (Fig. 5). The *FW300-53* genera include two genomes: one from Labrador Shelf sediment incubated with diesel and another from groundwater in Tennessee listed as an uncontaminated well nearby sites exposed to early nuclear research under the Manhattan Project [72].

### CuMMO Phylogeny

Interestingly, out of the 176 genomes analyzed in the *Porticoccaceae* family, the only genomes to encode particulate hydrocarbon monooxygenase (*pmoCAB*) belong to the *HB2-32-21, B045*, and *FW300_53* genera (Fig. 5). This suggests that the acquisition of *pmo* and the resulting metabolic function could be intertwined with the evolution of these three closely related genera. To further explore whether pmo is a potential driver of the *Porticoccaceae* evolution we assessed the frequency of genes further down the metabolic pathway for MCH consumption. We find genes involved in shunting MCH-derived carboxylic acids into central carbon metabolism to be uniquely abundant in *B045, HB2-32-21*, and *FW300_53* but largely absent from distant relatives in the *Porticoccaceae* family (Table S2). Within the *B045* genus, the amino acid identity of the *pmoA* subunit varies from 82-100%. Representatives from the *pmo* and *amo* enzyme class, called the CuMMO enzyme superfamily, can be found across the tree of life of bacteria and archaea and we sought to understand where the *pmo* from *B045* and its close relatives exists within the diversity of the CuMMO superfamily. Here, we analyze the phylogenetic relationship of all archaeal and bacterial *amo* and *pmo* sequences by using the alpha subunit as a phylogenetic marker similar to the method previously applied in other studies [59, 60]. By using B045-MAG *pmoA* sequences to query the National Center Biotechnology Information (NCBI) and Department of Energy Joint Genome Institute (DOE-JGI) databases we found more distantly related sequences to the novel *pmo* sequences, which originate from diverse environments including contaminated groundwater from mine drilling fluid, agricultural soil, brackish Black Sea water, and hydrothermal vents.

From our phylogenetic analysis, we observe all proteobacterial and Verrucomicrobia *amoA* and *pmoA* (including sequences that function on methane, ethane, butane, and ethylene) form a monophyletic clade which excludes B045-related *pmoA* sequences. Sequences related to B045-*pmo*, HB2-32-21-*pmo, FW300_53-pmo*, and the environmental relatives form a single monophyletic clade (Fig 6). Our phylogeny is largely consistent with a recently published tree on the CuMMO enzyme superfamily except for the placement of the Verrucomicrobia and NC10 clades, which in our tree branch with the proteobacterial sequences with high bootstrap support. In the previous study these clades branched closer to the gram-positive actinobacteria with poor bootstrap support [73]. Future research examining the expression of these genes at the RNA or protein level in cultures or environmental incubations, in tandem with extended geochemical analyses, may aid in extending our understanding of the function *B045 pmo* and the role it plays in the metabolism of the *B045* genera.

**Figure 6.**
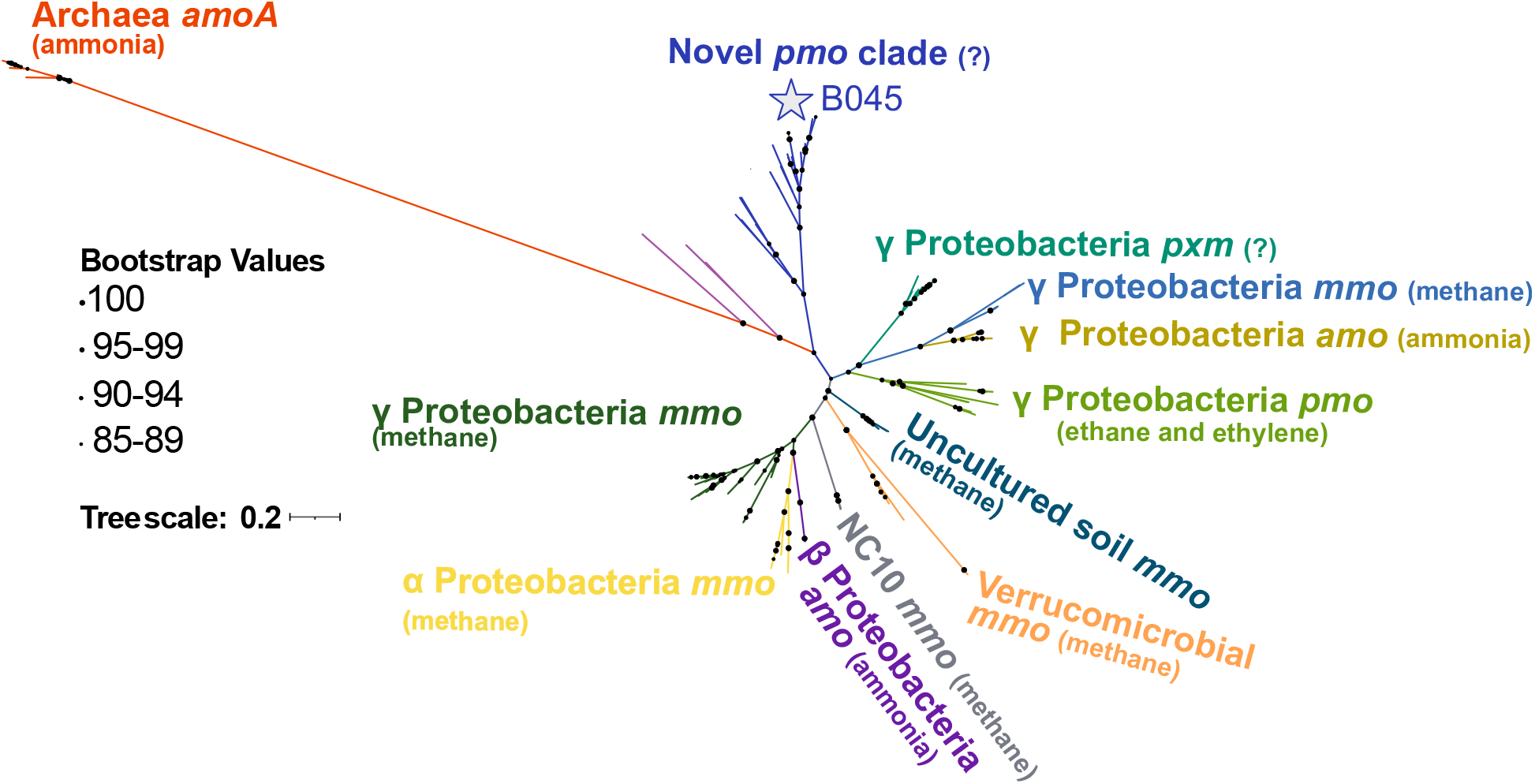
Phylogeny of *pmo*, *mmo*, and *amo* subunit A amino acid sequences. Maximum likelihood tree is drawn to scale, with branch lengths representing the number of substitutions per site. Bootstrap values below 50% are not shown. Each major clade is color coded for readability. Tree scale represents the number of substitutions per site. The B045 sequences are denoted by a star symbol.

### Conclusion

Our study provides the first genomic evidence for methyl-cycloalkane metabolism by the free-living novel *B045* genera within the *Porticoccaceae* family and further points to the environmental importance of these organisms and their metabolic activity. We have elucidated the potential pathways for MCH consumption which is the first environmental genomic study of methyl-cycloalkane consumption. Through the comparison of *Porticoccaceae* genomes and metagenomes, we show that the acquisition of the particulate hydrocarbon monooxygenase (*pHMOCAB*) may have been a key event in the evolution of hydrocarbon metabolism within the *B045* genera and the closely related *HB2-32-21* and *FW300_53* genera. The phylogenetic analysis of the CuMMO enzyme superfamily revealed the placement of the *pmo* from *B045* (and its close relatives) is separated from all other proteobacterial *amo/mmo/pmo* and it forms a novel monophyletic clade, providing insight into the evolutionary history of this important enzyme family.

## Data Availability

Metagenomes are available through NCBI in BioProject PRJNA897081.

## Acknowledgments

We thank Sasvath Ramachandran and Joshua Qin for their support in the bioinformatic analyses; the R/V Atlantis Captain and crew for support at sea. For bioinformatic analyses, this work used the Bridges-2 system, which is supported by NSF award number OAC-1928147 at the Pittsburgh Supercomputing Center (PSC). We thank David O’Neal and TJ Olesky for their assistance with bioinformatics optimization and support on the Bridges and Bridges-2 systems.

