## Supplementary Information for "Methylated Cycloalkanes Fuel a Novel Genera in the *Porticoccaceae* Family and Inform Substrate Affinity for a Unique Copper Membrane Monooxygenase"

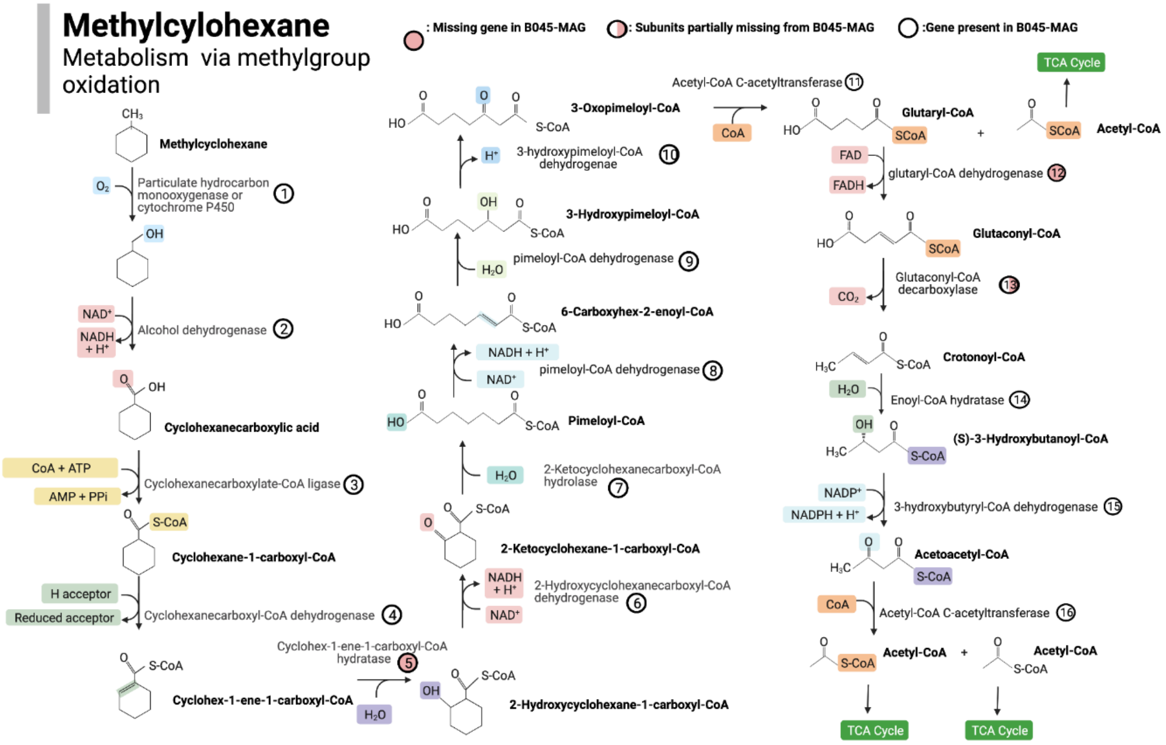


**Figure S1.** Hypothetical pathway for methylcyclohexane consumption via methyl group oxidation.

**Table S1.** KEGG Gene Identifiers used for Fig. S1. K0 ID's in bold are missing from B045-MAG.

| Metabolic Step | Enzyme Name | KO ID |
|---|---|---|
| 1 | Particulate hydrocarbon monooxygenase or cytochrome P450 | K10944 or K15486 |
| 2 | Alcohol dehydrogenase | K00138 |
| 3 | Cyclohexanecarboxylate-CoA ligase | K04116 |
| 4 | Cyclohexanecarboxyl-CoA dehydrogenase | K04117 |
| 5 | Cyclohex-1-ene-1-carboxyl-CoA hydratase | **K07534** |
| 6 | 2-hydroxycyclohexanecarboxyl-CoA dehydrogenase | K07535 |
| 7 | 2-ketocyclohexanecarboxyl-CoA hydrolase | K07536 |
| 8 | Pimeloyl-CoA dehydrogenase | K04118 |
| 9 | Pimeloyl-CoA dehydrogenase | ? |
| 10 | 3-hydroxypimeloyl-CoA dehydrogenase | K07516 |
| 11 | Acetyl-CoA acyltransferase | K00632 |
| 12 | glutaryl-CoA dehydrogenase | **K16173** |
| 13 | Glutaconyl-CoA decarboxylase | **K01615**, K20509, **K23351**, **K23352** |
| 14 | Enoyl-CoA hydratase | K01692 |
| 15 | 3-hydroxybutyryl-CoA dehydrogenase | **K00074** |
| 16 | Acetyl-CoA C-acetyltransferase | K00626 |

**Table S2.** Average number of copies of genes per phylogenetic group. Calculations were determined by gene copy number (from kofamscan, divided by number of genomes, which results in the metric # copies gene/genome)

| Metabolic Step | Enzyme Name | KO ID | *B045* Genus | *HB2-32-21* Genus | *FW300_53* Genus | Distant Relatives *Porticoccaceae* |
|---|---|---|---|---|---|---|
| 1 | Particulate Hydrocarbon Monooxygenase | K10944 | 1 | 1 | 1.2 | 0 |
| 2 | Cyclohexanol Dehydrogenase | K19960 | 0.125 | 0 | 0 | 0 |
| 3 | Cyclohexanone Monooxygenase | K00499 | 1.5 | 1 | 2 | 2.8 |
| 4 | Gluconolactonase | K03379 | 5.625 | 8 | 7.4 | 1 |
| | Epsilon-Lactone Hydrolase | K14731 | 5 | 4 | 7.2 | 0.4 |
| 5 | Alcohol Dehydrogenase | K13954 | 0.75 | 0 | 1 | 0.2 |
| 6 | Aldehyde Dehydrogenase | K00138 | 1 | 1.5 | 1.8 | 0 |
| 8 | Acyl-CoA Dehydrogenase | K00249 | 15.5 | 10.5 | 15.4 | 2.6 |
| | enoyl-CoA hydratase | K01692 | 5.25 | 3 | 7.6 | 3.4 |
